# Losses of *TMC* and *CIB* neurosensory genes in schizophoran flies at the PETM

**DOI:** 10.64898/2026.07.31.741574

**Authors:** Haley V. Reeves, Cal Baker, Aleicea Rodriguez, John M. Logsdon, Albert Erives

**Affiliations:** Department of Biology, University of Iowa, Iowa City, IA, 52242-1324 USA

**Keywords:** Transmembrane channels (TMCs), calcium- and integrin-binding (CIB) proteins, Schizophoran radiation, thermal nociception, Paleocene-Eocene thermal maximum (PETM)

## Abstract

The K–Pg extinction ended the world of non-avian dinosaurs 66 Mya and ushered in the Age of Mammals. During the ∼10 My of recovery, Schizophora, an immensely successful group of flies, appeared, flourished and diversified. This large radiation produced over half of all dipteran families (∼78/150), including the family for *Drosophila*, the model genetic powerhouse. In the context of this evolutionary radiation, we investigate the loss of two of three highly conserved, neurosensory transmembrane channel (TMC) genes. Here, we show that these genes were separately lost across multiple diverging schizophoran lineages of the early Paleogene, suggesting a powerful environmental driver was involved. We also show that unlinked genes encoding the calcium- and integrin-binding (CIB) subunits of the missing TMC complexes were lost during the same time frame. Because the lost genes encode complexes involved in thermal nociception, we propose the external driver was likely the Paleocene-Eocene thermal maximum (PETM), a 200 ky interval of elevated global temperatures occurring 56 Mya, slightly before the earliest schizophoran fossil from 53 Mya. These results suggest that gene loss may have been adaptive for most schizophoran lineages to emerge past the hothouse Earth of the PETM.

## Introduction

The Cretaceous-Paleogene (K–Pg) extinction marks the beginning of the Cenozoic Era with a thin layer of clay sediment enriched in iridium due to the Chicxulub impactor event 66.043 Mya that caused extinction of ∼73% of the world’s species (1). The overturning of global flora and fauna, recorded geologically, allowed new evolutionary radiations of placental mammals, birds, insects, and angiosperms. These post K–Pg radiations are also detectable in phylogenetic analyses as bursts of diversification. While the K–Pg extinction event is not the largest of the traditional “Big Five” extinction events in Earth’s history, it is the most recent one and, as such, the most accessible to phylogenetic analysis of its aftermath.

Here we detail a series of discoveries that newly link the post K–Pg world of the early Paleogene with two established areas of study: (1) a pair of interacting gene families associated with animal sensory biology and (2) the evolutionary radiation of Schizophora, the largest diversification of animals in the Cenozoic. Schizophora are a section of true flies that evolved the “ptilinum”, a specialized, inflatable and retractable sac the adult uses to break out of the puparium, leaving a visible frontal suture that is the basis for their name (Schizophora = “split-bearers”). Schizophoran flies are missing from the Cretaceous, first appeared in the early Paleogene, and gave rise to 78 extant families. Because of the rapidity and immensity of this evolutionary radiation, the relationships amongst schizophoran lineages have been a long-standing and vexing problem in evolutionary studies (2, 3)—consequential in motivating development of phylogenetic systematics and its use of synapomorphic characters over homoplasies (4).

We investigated two functionally integrated gene families. The first family encodes the transmembrane channels (TMC), which have been of interest due to the involvement of a subset of paralogs in the mechanotransduction channel complexes underlying vertebrate hearing and vestibular sensory systems (5–9). Other TMC paralogs have been implicated in other sensory modalities in both vertebrates and insects including sensation of texture, saltiness, noxious heat, pain, and itching (10–17). The second set of genes encodes calcium- and integrin-binding (CIB) proteins, which are required intracellular subunits in different types of TMC complexes (9, 10, 18–23). Here, we first show that most invertebrate eumetazoans are characterized by three paralogs, which are further duplicated in vertebrates. Second, we show that most schizophoran flies, including *Drosophila*, are missing up to two of the three genes from each family for a total of four missing genes in flies closely related to *Drosophila*. To identify when in the ancestry of *Drosophila* these genes were lost, we meticulously curated these genes from hundreds of genomes and conducted phylogenetic analyses. In doing so, we realized that these gene losses were more complex and significant than originally envisioned.

We find that the timing of gene losses within the TMC and CIB families of schizophoran flies related to *Drosophila* cannot be explained by single gene loss events, but that an external environmental driver must have selected for flies with these gene losses. Because homologs of some of these lost genes have been implicated in thermal nociception, *i.e.*, the sensation of noxious levels of heat stress (13, 14), we tentatively propose here that the Paleocene-Eocene thermal maximum (PETM) was that external driver. The extreme hothouse Earth of the PETM occurred well after the Schizophoran radiation was underway but ended just before the earliest fossils of schizophoran material. Thus, these findings may be relevant to understanding the types of genetic resiliency available to some animals during significant climatic warming trends. It also suggests that gene loss may have been adaptive, allowing some cyclorrhaphan fly lineages to flourish and continue the evolutionary radiation of Schizophora past the PETM.

## Results

### Loss of TMC genes in early Schizophora

This study originated as a follow-up investigation regarding the timing of the loss of two highly conserved genes in the ancestry of *Drosophila melanogaster*: *Tmc487* and *Tmc56* (Fig. 1A). Because of these absences and our prior study’s focus on early branching metazoans, the single *Drosophila* TMC gene was excluded (24). Figure 1A is an updated phylogenetic analysis of the metazoan TMC family by maximum likelihood (ML) estimation, now including *Drosophila Tmc123* and additional, relevant, schizophoran TMC genes (see red asterisks in Fig. 1A). This new tree demonstrates that Eumetazoa is characterized by three TMC genes (*Tmc123*, *Tmc487*, and *Tmc56*), which further duplicated into *TMC1*–*TMC8* in jawed vertebrates. Unlike most eumetazoans, the schizophoran fly *Drosophila melanogaster* retains only the *Tmc123* gene, known as *Tmc*, despite the lost genes being highly conserved across Metazoa (Fig. 1A). We show here that the timing of these losses occurred multiple times during the early evolutionary radiation of Schizophora.

**Figure 1.**
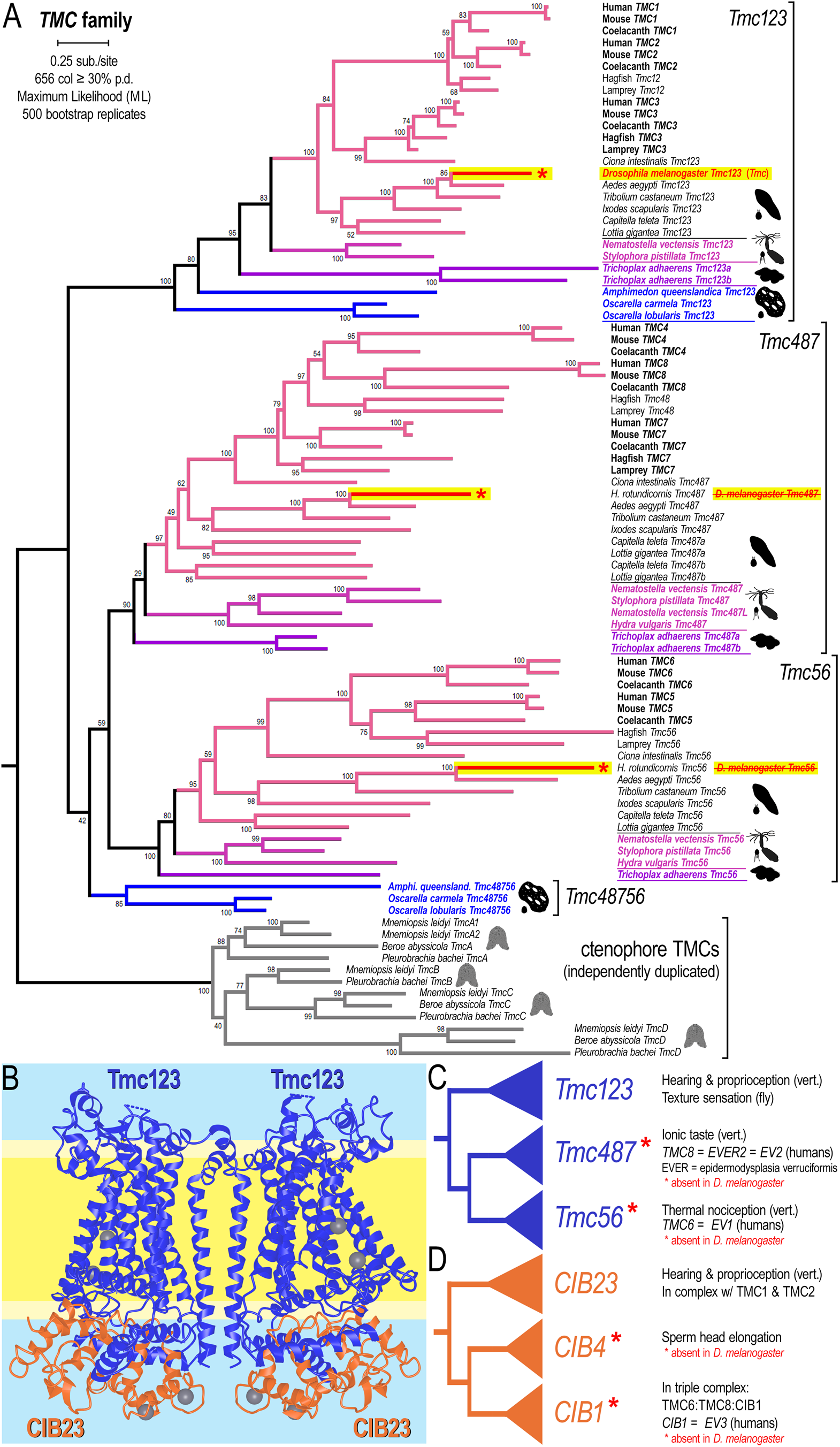
Loss of *Tmc487* and *Tmc56* in the schizophoran ancestry of *Drosophila melanogaster*. **(A)** Phylogenetic analysis of the metazoan TMC family by maximum likelihood (ML) estimation shows that Eumetazoa (Placozoa + Cnidaria + Bilateria) is characterized by three TMC genes (*Tmc123*, *Tmc487*, and *Tmc56*), which further duplicated into *TMC1*–*TMC8* in jawed vertebrates. Unlike most eumetazoans, the schizophoran fly *Drosophila melanogaster* retains only the *Tmc123* gene (*Tmc*) as it is missing *Tmc487* and *Tmc56* (lineages with red asterisks). In place of *Tmc487* and *Tmc56* from *Drosophila*, this tree features another fly, *Heteromyza rotundicornis* (see Fig. 2B, one of the few schizophorans with the full ancestral eumetazoan repertoire. The evolutionary radiation of Schizophora was the largest radiation of animals of the Cenozoic Era. The immensity and rapidity of this radiation have obscured internal relationships amongst schizophoran lineages. The 656 alignment columns (“col”) with ≥ 30% partial data (“p.d.”) were used to construct this tree. See Methods for details. **(B)** Structure of the native Tmc123 channel complex from *C. elegans* (TMC-1) is composed of a TMC dimer (blue transmembrane subunits) and two calcium- and integrin-binding subunits (orange intracellular subunits with gray Ca^2+^ atoms). *C. elegans* CALM-1 is referred to here as CIB23 for its orthology to vertebrate CIB2 + CIB3 subunits, like our TMC naming scheme. **(C,D)** The tri-partite topology of the eumetazoan TMC tree (C, blue tree) is recapitulated in the CIB tree (see Fig. S1), which is composed of *CIB23*, *CIB4* and *CIB1* (D, orange tree). *Drosophila melanogaster* is also missing *CIB4* and *CIB1* in addition to *Tmc487* and *Tmc56* (genes with red asterisks). Altogether these genes encode channel subunits involved in diverse sensory functions (some listed on the right in C and D), including thermal nociception. Human *TMC6* (*EV1*), *TMC8* (*EV2*), and *CIB1* (*EV3*) underlie genetic causes of epidermodysplasia verruciformis (EVER).

### Loss of CIB genes alongside TMC losses

The structures of the native *Tmc123* channel complexes from *C. elegans* (TMC-1 and TMC-2, from a pair of nematode-specific *Tmc123* duplications) were recently determined (22, 23). These complexes are composed of TMC dimers (blue transmembrane subunits in Fig. 1B) and two intracellular calcium- and integrin-binding subunits (called CALM-1 in *C. elegans*; orange subunits with gray Ca2+ atoms in Fig. 1B). Furthermore, *CIB2* and *CIB3* are known deafness genes that encode intracellular EF-hand subunits of the TMC1 and TMC2 mechanotransducing complexes that underlie hearing and vestibular sensory structures of the inner ear (6). Similarly, TMC6 and TMC8 form a heteromeric complex with CIB1 (21). In humans, *TMC6* (*EV1*), *TMC8* (*EV2*), and *CIB1* (*EV3*) encode subunits of a heteromeric complex and correspond to the three main loci mutated in epidermodysplasia verruciformis (EVER) (25–27). Thus, both genetic and biochemical lines of evidence have been found connecting specific TMC paralogs with specific CIB paralogs. Here, we add a third line of evidence, which is the phylogenetic co-evolution (rates and losses) between specific pairs of TMC and CIB paralogs.

To see if a subset of CIB paralogs is also lost in Schizophora, we conducted phylogenetic analyses of an extensive set of curated CIB genes from eumetazoans with an emphasis on cyclorrhaphan flies (see Methods). We find that the tri-partite topology of the eumetazoan TMC tree (24) (Fig. 1C, blue tree) is recapitulated in the CIB tree (Fig. 1D cartoon, and Fig. S1 for a full tree), which is composed of the paralogy clades *CIB23*, *CIB4* and *CIB1* (Fig. 1D, orange tree). Moreover, we find that *Drosophila melanogaster* and related schizophoran flies are missing *CIB4* and *CIB1* in addition to *Tmc487* and *Tmc56* (genes with red asterisks in Fig. 1A and documented in more detail in later figures). Altogether these genes encode channel subunits involved in diverse sensory functions (some listed on the right in Fig. 1C–1D), including thermal nociception (13, 14).

### Many separate losses of CIBs and TMCs

We find that the highly conserved *CIB1* gene is missing from the genome assemblies of 26 of the 78 schizophoran families for which genome assemblies are available (∼31 families). These 26 families are distributed in 9 different superfamilies. This gene is also missing in the Syrphinae, but not the Eristalinae subfamily of Syrphidae, a basal cyclorrhaphan family, which suggests that some unknown factor specifically predisposed cyclorrhaphan lineages for these gene losses, unlike most other animals. Within Schizophora, *CIB1* is present in all known heleomyzids with genomic data (*n* = 4 species from two subfamilies), one of two available diopsids, and in some families of the Tephritoidea superfamily (Fig. 2A). *CIB1* is notably absent in the Tephritidae family, for which genomes from multiple genera have been sequenced and assembled (40 species from 19 tephritid genera to date). For this study, we also confirmed all gene absences at the level of sequencing reads, which we also used to correct mis-assembled regions.

**Figure 2.**
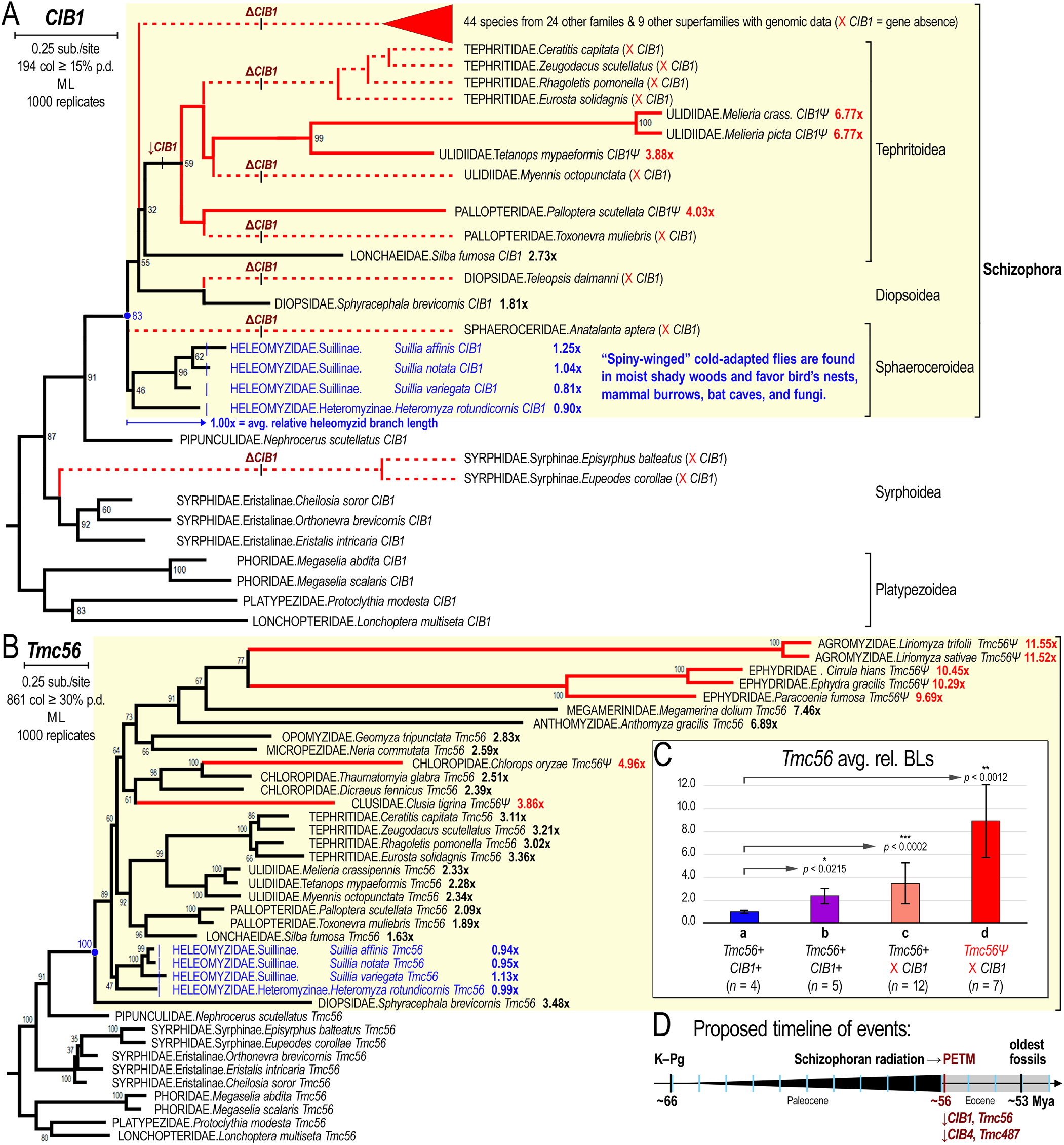
*CIB1* and *Tmc56* were separately lost multiple times within Schizophora after the schizophoran radiation. **(A)** Shown is the cyclorrhaphan *CIB1* tree with Schizophora highlighted (yellow box). The highly conserved *CIB1* gene is missing from the genome assemblies of 26 of the 78 schizophoran families for which genome assemblies are available (∼31 different families). These 26 families are distributed in 9 different superfamilies. This gene is missing also in the Syrphinae, but not the Eristalinae subfamily of Syrphidae, a basal cyclorrhaphan family. Within Schizophora, *CIB1* is present in all known heleomyzids with genomic data, one of two available diopsids, and in some families of Tephritoidea, but notably absent in Tephritidae, for which genomes from multiple genera have been sequenced (40 species from 19 genera to date). *CIB1* is absent in all other available genome assemblies within Schizophora. Lineages with missing genes from flies analyzed in this study are shown in dotted red lines (“X *CIB1*”). Lineages with pseudogenes are indicated with a “*Ψ*” suffix in the gene name. Pseudogenes are characterized by missing splice sites, open reading reading frame slips, and/or truncated genes (missing exons) in both the genome assembly and across all available sequencing reads. Because all flies within Tephritoidea are either missing *CIB1* (dark red “Δ*CIB1*” along dotted red lineages) or possess a pseudogenized *CIB1Ψ* (solid red lineages), loss of selection can be inferred to have occurred early in the radiation of Tephritoidea (dark red “↓*CIB1*”). To characterize the pace of pseudogenization and gene loss within Schizophora, relative branch lengths were computed by normalizing distances by the average heleomyzid branch length, which we define as 1.0 (blue dotted “1.00” line). Branch lengths are measured from the base of Schizophora (blue node) to the lineage tips and normalized by the average absolute branch length for heleomyzids. Relative branch lengths are shown to the right of the taxonomic names. **(B)** Shown is the cyclorrhaphan *Tmc56* tree with Schizophora highlighted (yellow box). The *Tmc56* tree shows that heleomyzids are the least derived schizophoran lineages. *Tmc56Ψ* pseudogenes are indicated in red as in panel A, but lineages with missing genes are not shown. **(C)** Average Tmc56 helomyzid- normalized branch lengths (“BLs”) are shown for four different, mutually exclusive data sets: (a) heleomyzids, (b) non-heleomyzids with *CIB1* genes, (c) (non-heleomyzid) flies missing *CIB1*, and (d) flies with *Tmc56Ψ* pseudogenes, all of which are non-heleomyzids missing *CIB1*. In comparison to heleomyzids (set a), all other schizophorans have significantly different BLs (asterisked *p*-values from one-tailed student *t-*tests under the assumption of unequal variances). **(D)** Because the multiple, separate losses of *CIB1*, *Tmc56*, *CIB4*, and *Tmc487* constitute a dramatic series of events requiring an external or environmental driver, and because these genes encode interacting subunits involved in thermal nociception among other sensory functions in vertebrates, we tentatively suggest that these losses are associated with climatic events during and around the PETM. The PETM was a brief 200 ky interval during which global temperatures increased ∼5°C to produce a hothouse Earth, the warmest known interval in the last 66 My.

In support of the robustness of gene detection methods used in this study, we also identified pseudogenized copies of some of these genes, particularly in clades known to harbor both missing and intact genes. For example, we find lineages within the superfamily Tephritoidea that contain intact *CIB1* genes, ones that contain *CIB1Ψ* pseudogenes (solid red lines in Fig. 2A) or complete absences (dotted red lines in Fig. 2A). These pseudogenes are characterized by missing splice sites, reading frame slips, and/or truncated genes (missing internal coding exons) in both the genome assembly and across all available sequencing reads. We are thus confident we can distinguish fast-evolving protein-coding genes from pseudogenized, non-functional remnants of them. We further find that empirically we cannot detect intact genes or pseudogenes that are ∼12.3x more divergent than the least derived schizophoran lineages (explained below).

Genome assemblies from flies within Tephritoidea have *CIB1* genes that are fast-evolving (*n* = 1 from Longchaeidae), pseudogenized (*n* = 4, *CIB1Ψ* solid red lineages from Ulidiidae and Pallopteridae in Fig. 2A), or missing (all tephritids, and some lineages from Ulidiidae and Pallopteridae, dark red “`Δ`*CIB1*” along dotted red lineages in Fig. 2A). Thus, a loss or reduction of selection can be inferred to have occurred early in the radiation of Tephritoidea (e.g., dark red “↓*CIB1*” in Fig. 2A). In total, there are at least three complete losses of *CIB1* within Tephritoidea. In addition, there are separate *CIB1* losses in Diopsidae, Sphaeroceridae, and a hypothetical stem lineage leading to the remaining bulk of extant schizophorans. These three losses of *CIB1* cannot be equated with the separate losses within Tephritoidea, whose monophyly and composition is highly supported. But even if the presumed sphaerocerid loss of *CIB1* was equated with a hypothetical single loss leading to the bulk of Schizophora, there would be at least six separate losses of *CIB1* across Cyclorrhapha (including the absence from the Syrphinae subfamily of Syrphidae; see Fig. 2A and Table 1).

**Table 1.** Conservative TMC and CIB gene loss estimates across Cyclorrhapha. Conservatively, there are a minimum of 18–19 separate gene losses in Schizophora, and 19–20 separate gene losses in Cyclorrhapha. See Fig. S2 for an indexed count of gene loss characters mapped onto a cladogram.

| Gene | Pseudogenes counted as genes |  | Pseudogenes counted as losses |  |
| --- | --- | --- | --- | --- |
|  | Schizophora | Cyclorrhapha | Schizophora | Cyclorrhapha |
| *CIB1* | 5 | 1 | 3 | 1 |
| *Tmc56* | 4 | 0 | 5 | 0 |
| *CIB4* | 4 | 0 | 4 | 0 |
| *Tmc487* | 6 | 0 | 6 | 0 |
| **Total losses** | **19** | **1** | **18** | **1** |

No clear *CIB1Ψ* pseudogenes were found outside of Tephritoidea, but at least four separate *Tmc56Ψ* pseudogenizations were found in schizophoran genomes missing *CIB1* (red lineages leading to Agromyzidae, Ephydridae, Chloropidae, and Clusidae in Fig. 2B). Altogether, we find that there had to be at least nine separate losses of *CIB1* and *Tmc56* across Schizophora when estimated conservatively in a maximally plausible polytomous tree (see indexed losses in Fig. S2). The use of a maximally polytomous tree has two functions: (*i*) it sidesteps a need to know the exact topology in the deep parts of the schizophoran tree, and (*ii*) it allows us to minimize the number of losses and get an absolute lower bound. Similarly, we characterized an additional ten separate losses for *CIB4* and *Tmc487* (Table 1 and Fig. S2). Finally, this absolute minimum number of losses in Schizophora is not greatly affected by counting pseudogenes as genes (*n* = 19 separate losses) or as losses (*n* = 18 separate losses). Thus, each of the four genes was lost a minimum of 4–6 times and likely more than this during the schizophoran radiation.

To characterize the pace of CIB and TMC pseudogenization and gene loss within Schizophora, relative branch lengths (“BLs”, representing relative rates of evolution) were computed by normalizing distances by the average heleomyzid branch length, which we define as 1 (see blue dotted lines “1.00” lines in Fig. 2A–2B). Branch lengths are measured from the base of Schizophora (blue nodes in Fig. 2A–2B) to the lineage tips and normalized by the average absolute branch length for heleomyzids. Relative branch lengths are shown to the right of the taxonomic names in Figure 2 trees.

Like the *CIB1* tree, the *Tmc56* tree shows that heleomyzids are the least derived schizophoran lineages that harbor remnants of these genes (Fig. 2B). Furthermore, we can also find distinct *Tmc56Ψ* pseudogenes (indicated in red in Fig. 2B). In Figure 2C, average Tmc56 helomyzid-normalized branch lengths are shown for four mutually exclusive sets of schizophoran flies: (a) heleomyzids, which have all been found to have short-branched *CIB1* genes; (b) non-heleomyzids with *CIB1* genes, which have been found to be long-branched; (c) non-heleomyzids, which are missing *CIB1*; and (d) non-heleomyzids with *Tmc56Ψ* pseudogenes, all of which are also missing *CIB1*. In comparison to heleomyzids (set A), all other schizophorans have significantly different BLs, thus supporting the idea that heleomyzids possess the least derived versions of these genes. This also shows that fast-evolving *Tmc56* genes in Schizophora are correlated with faster evolutionary rates of their *CIB1* genes. Below we will discuss how these pseudogenizations and gene losses make the most sense in the context of the Paleogene greenhouse and the PETM (Fig. 2D).

A heat map of relative branch lengths and gene losses for all analyzed genes demonstrates the contrast of most schizophoran lineages with the heleomyzids (Fig. 3A). We find that heleomyzid-normalized branch lengths for all three pairs of genes *Tmc123-CIB23*, *Tmc487-CIB4*, and *Tmc56-CIB1* have branch lengths that are all correlated to one another within each lineage despite different genes evolving at different rates (Pearson’s *r* ranges from 0.50–0.80, see Fig. 3B). A comparison of *Tmc123-CIB4* coevolution (*n* = 32), a mismatched pairing, is poorly correlated (*r* = 0.209) unlike the high correlation for the same *Tmc123* values paired with *CIB23* (*r* = 0.821, see Fig. S3).

**Figure 3.**
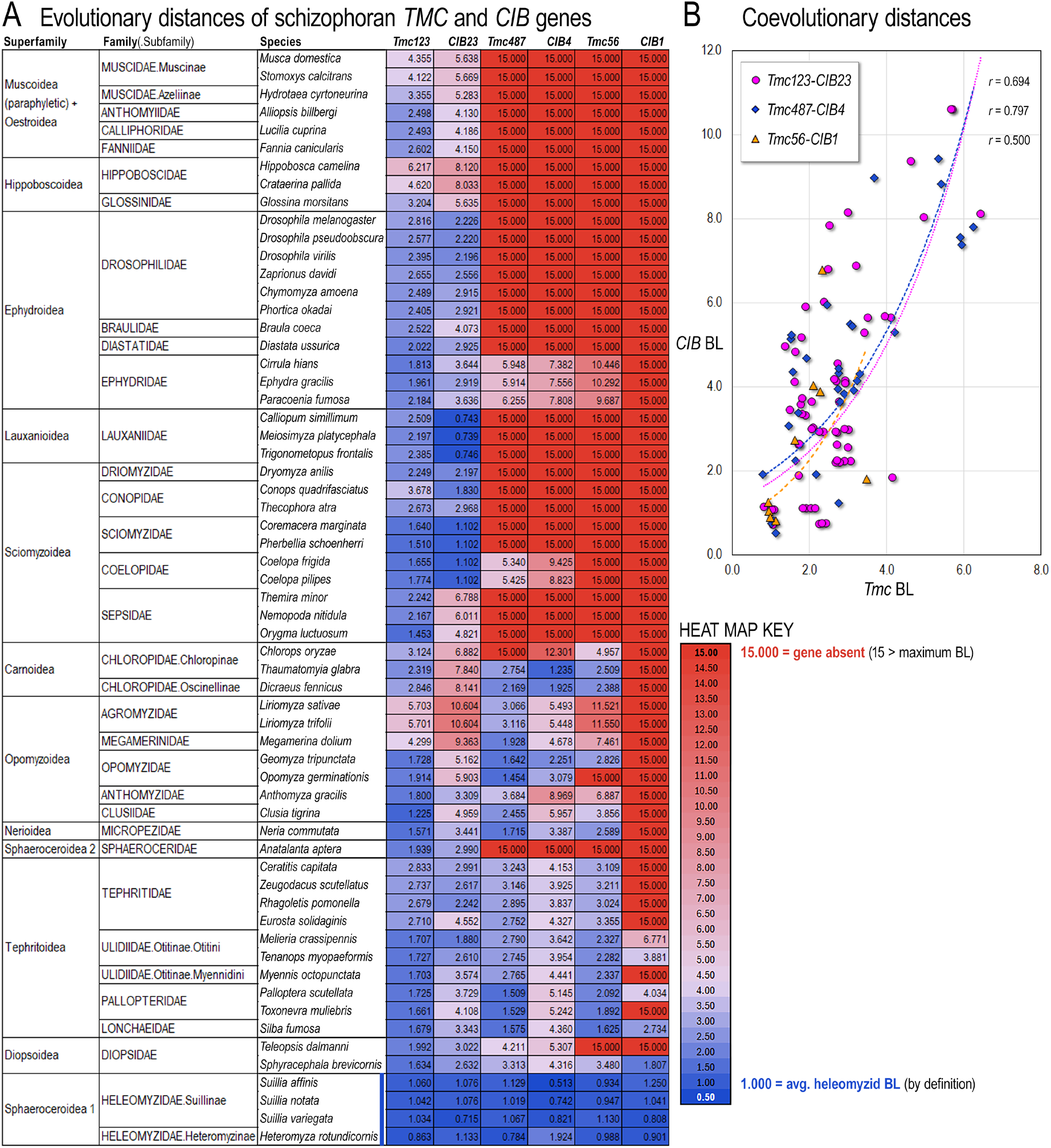
Evolutionary rates and losses of TMC paralogs are correlated to specific CIB subunits. **(A)** Shown is a heat map of relative branch lengths (BL) for all TMC and CIB schizophoran genes. A relative BL of 15.00 (red ceiling in the key) was chosen to represent complete gene absence, as no measured BL was greater than ∼12.3. A relative branch length of 0.50 (blue floor in the key) was chosen to represent the shortest branch length, as the smallest BL was 0.513. The average heleomyzid BL was defined to equal 1. **(B)** Scatter plots between BLs of matched pairs of TMC and CIB paralogs show that rates are moderately (Pearson’s *r* = 0.50) to highly correlated (0.80) unlike a mismatched pairing (*r* = 0.21, see Fig. S3). Trendlines are exponential fits to the three data sets, *Tmc123-CIB23* (magenta short-spaced dotted line), *Tmc487-CIB4* (blue medium-spaced dotted line), and *Tmc56- CIB1* (orange wide-spaced dotted line).

These four gene losses from the TMC and CIB families are unusual and call out for an explanation. Excluding Schizophora, we find all three TMC and all three CIB genes in all major insect orders of Holometabola (specifically non-schizophoran Diptera, Lepidoptera, Trichoptera, Coleoptera, and Hymenoptera, *e.g.*, see the CIB23 tree in Fig. S1). Representative trees for *CIB4*, *Tmc487*, and *CIB23* genes within Cyclorrhapha are shown in Figs. S4–S6, respectively, and for the *Tmc123* tree in Fig. 4.

**Figure 4.**
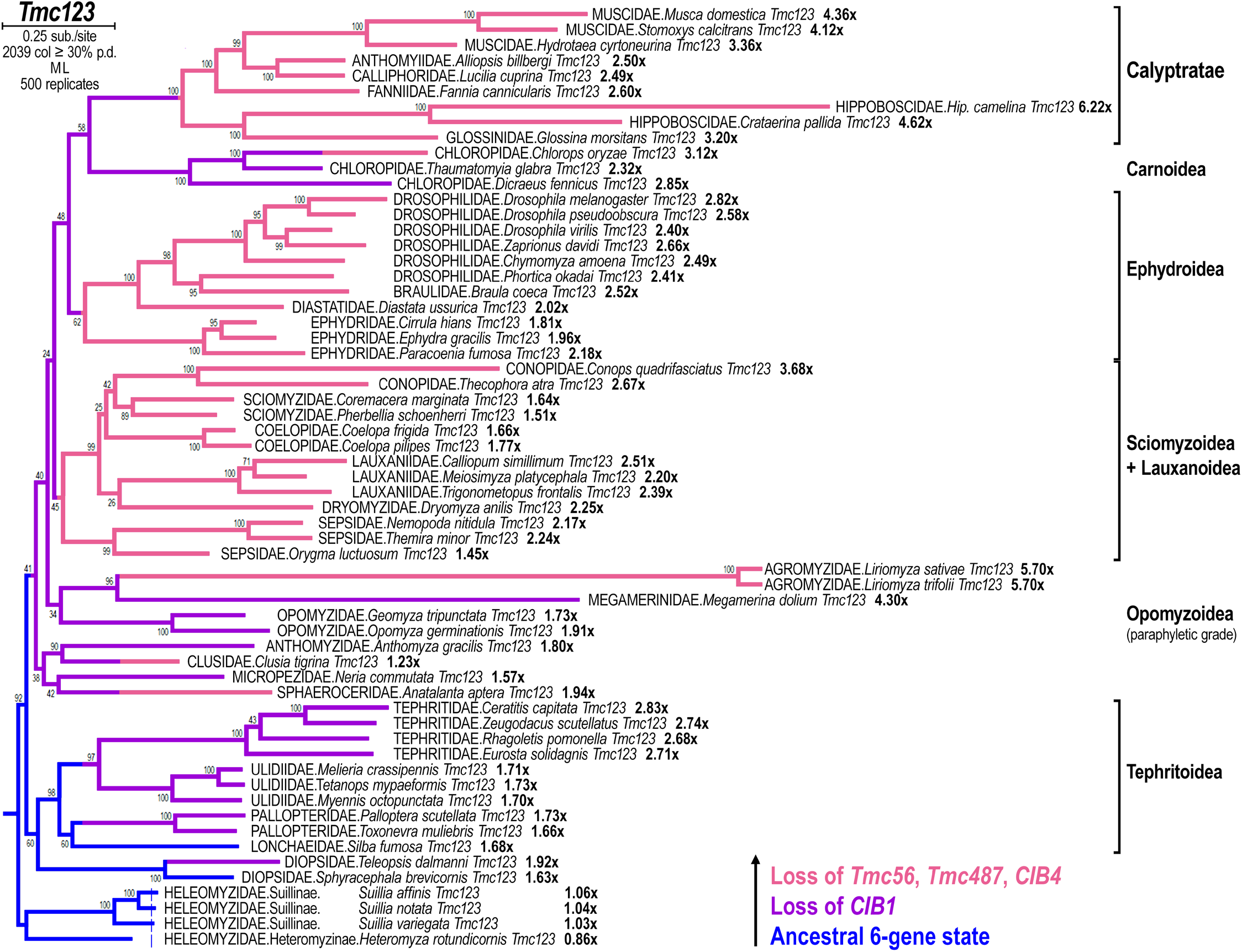
Basal schizophorans have ancestral repertoires of *TMC* and *CIB* genes. Phylogenetic analysis of schizophoran *Tmc123* genes, which have not been found missing in any lineage, places the lineages with the most complete *TMC* and *CIB* repertoires at the base of Schizophora. After the basal heleomyzids, there is a super clade composed of Tephritoidea and Diopsidae in one clade and all other schizophorans in a sister clade. While there are some fast-evolving (long-branched) genes, none of these have clear signatures of pseudogenization. This tree is color-coded by the pattern of gene losses (see key in lower right-hand corner with arrow corresponding to direction of time and nested pattern of losses).

### PETM relevance to TMC and CIB losses

One early motivation for pursuing these investigations was the possibility that relationships amongst schizophoran lineages would be resolved by a consistent pattern of a few gene losses. However, the data were overwhelmingly clear that single gene loss events were untenable as an explanation worth investigating with more complicated analyses. For example, the loss of *CIB1* cannot be constrained to a single event in the Schizophoran radiation without overturning well-established schizophoran relationships (2, 3). In particular, the multiple losses within Tephritoidea cannot be equated to a single loss with those outside the superfamily. Thus, to explain the multiple losses of *CIB1*, *Tmc56*, *CIB4*, and *Tmc487*, these dramatic events require some external or environmental driver. Because these genes encode interacting subunits involved in thermal nociception and other stress responses in vertebrates (13, 14), we tentatively propose that these losses are associated with climatic events during and around the PETM (see Fig. 2D). The PETM was a brief 200 ky interval during which global temperatures increased ∼5^°^C to produce a hothouse Earth, the warmest known interval in the last 66 My (28). Furthermore, the PETM is known to have resulted in widespread disruption of terrestrial ecosystems (29). These results may illustrate a new way in which the PETM and other parts of Earth’s history may be read genomically.

### Basal schizophorans retained all genes

Phylogenetic analysis of schizophoran *Tmc123* genes, which we have not found missing in any lineage, places the clades with the most complete *TMC* and *CIB* repertoires at the base of Schizophora. After the basal heleomyzids, there is a super clade composed of Tephritoidea and Diopsidae in one clade, and all other schizophorans in a sister clade. While there are some fast-evolving lineages, none of these have clear signatures of pseudogenization (see color-coded pattern of gene losses in Fig. 4.)

The phylogeny of Schizophora based on Tmc123 recovers the main monophyletic clades of (1) Calyptratae (the superfamilies of Muscoidea + Oestroidea + Hippoboscoidea), (2) Ephydroidea, (3) a clade composed of Sciomyzoidea + Lauxanoidea, (4) a clade composed of Tephritoidea + Diopsidae, and (5) the basal branching family Heleomyzidae (see named, bracket clades in Fig. 4). Carnoidea is represented only by Chloropidae, so its weak association as a sister clade to Calyptratae needs testing with lineages from other families. Last, Opomyzoidea appears as a paraphyletic basal grade that emerges as the sister clade to the clade composed of Tephritoidea + Diopsidae, consistent with modern systematics rejecting monophyly for this superfamily (30). The bulk of Schizophora characterized by the highest extent of TMC and CIB gene losses emerges out of this basal opomyzoid grade (see *Tmc123* tree color-coded by the general pattern of gene loss in Fig. 4). Finally, we also find a shared seven amino acid deletion associated with a phase change in intron 4 of *Tmc123*, which unites Pipunculidae with Schizophora to the exclusion of Syrphoidea, supporting the idea that Syrphoidea (Syrphidae + Pipunculidae) is not monophyletic (see Fig. S7).

This study adds to the growing body of novel phylogenetic results that eschew large data sets of single-copy (*i.e.*, unduplicated) genes in favor of paralogous gene families(24, 31, 32). For example, the gene duplication that resulted in *Tmc123* + *Tmc48756* unites Porifera with Eumetazoa in a proposed “Benthozoa” (24), while the multiple independent duplications of TMCs in the ancestry of ctenophores simultaneously excludes them from “Benthozoa” (Fig. 1A). Phylogenies of paralogous gene families also produce results that are unattainable with single-copy genes. For example, the phylogenetic analyses of eukaryotic core histones inherently result in trees that split the stem-eukaryotic lineage at the duplication points, which further suggest the evolutionary steps taken to arrive at linear chromosomes (31, 32). In contrast, single-copy genes of extant organisms are only able to define stem-lineages in deep time. Furthermore, gene duplications frequently result in neo-functionalizations, which better comport with synapomorphies of new clades and/or major transitions than do single-copy genes evolving at clock-like rates. Paralogous gene families like the TMCs, which have been adapted for different sensory modalities, or which have been partially lost in specific clades, will be intimately connected to the range of metazoan morphologies and behaviors.

## Discussion

### A PETM culling of early Schizophora

Here we showed that four genes encoding subunits of transmembrane complexes and their intracellular calcium binding subunits were all lost across the radiating schizophoran tree during the early Paleogene. The earliest accepted non-trace fossil of a schizophoran fly is from a lauxaniid in Baltic amber dating to 53 Mya after the PETM had concluded (33). As a lauxaniid is already from a well-established family from an unranked clade composed of the superfamilies of Sciomyzoidea and Lauxanoidea, the main lineages of Schizophora were already in place soon after the PETM (see Fig. 2D). So, if the proposed interpretation of these concerted gene losses is correct in their connection to the PETM, it would suggest that the early radiation of Schizophora in the Paleogene underwent bottlenecks at the PETM, such that primarily lineages with these gene losses came through and survived past the hothouse Earth of the PETM.

Given multiple, separate losses of the same set of four genes across different well-established parts of the schizophoran tree, it is likely there were selective benefits specifically in these fly lineages, but, for yet unknown reasons, not in other insects. In schizophorans, these genes possibly encoded a thermally induced repressive safety switch that needlessly prevented these flies from thriving in the new hot environment. While thermally regulated pathways are critical for coping with both daily temperature fluctuations and extreme temperature events, such as heat waves and cold snaps, some thermosensing circuits may have become maladaptive during the 200 ky hothouse of the PETM. Furthermore, the duration of the PETM onset (carbon isotope excursion onset) has been estimated to have taken a mere 1–6 ky (34–37). This geologically instantaneous warming compounded the existing greenhouse conditions of the early Paleogene (38).

The separate gene losses across most of the lineages of the early schizophoran radiation contrasts with the basal schizophoran family of Heleomyzidae, which we have found to carry the full complement of genes in at least two subfamilies (Suillinae and Heteromyzinae). Extant heleomyzids are noted as cold-adapted flies (39), favoring: cool dark woods (40); mammal burrows (41), which have fewer temperature extremes; and bat caves (42). The wombat flies of the heleomyzid Borboroidini tribe are found around the extensive burrows of wombats (41), which are among the largest burrowing mammals, are mainly crepuscular and nocturnal, and have diminished capacities to regulate body temperature during the day outside their burrows (43). Given these preferred habitats, refugial ancestors of modern heleomyzids may have been spared the effects of a geologically brief PETM culling and thus survived as a basal ancestral clade.

The peripheral nervous system of insects has several types of thermoreceptor neurons that send afferents to the CNS to influence behaviors such as thermotaxis and heat avoidance as well as associative memory (44, 45). These behaviors are guided by *at least* four classes of thermosensitive neurons that respond to innocuous cooling or heating, or noxious heat or cold (44). TRP channels have been found to be important thermoreceptors of sensory neurons mediating hot sensing and sensory neurons mediating sensing of innocuous cooling in the *Drosophila* antenna (46, 47), while ionotropic receptors have been found to mediate larval transition from early warm-sensing to late cool-sensing in *Drosophila* larvae (48). Thus, much comparative work using *Drosophila*, heleomyzids, and basal cyclorrhaphan lineages will be required to model how various thermosensory neurons may have functioned under different climatic conditions.

Citing observations on the behavior and anatomy of a calyptrate (the blowfly *Calliphora vomitoria*), Darwin famously wrote that the “*mental faculties of the Diptera are probably higher than in most other insects, in accordance with their highly developed nervous system.*” (49) This is particularly true of schizophoran flies, which exhibit some of the most complex behaviors, including aerial acrobatics controlled by advanced integration of chemical and visual avionics (50–53). Thus, these results raise the question as to whether early schizophorans may have wrested behavioral control from basic sensory-driven reflexes in favor of higher cognitive processing involving more broad-based sensory integration and associative memory.

## Methods

### Gene curation

To curate the TMC and CIB genes and correct errors inherent to many different approaches to *de novo* gene prediction, we curated most genes from genomic assemblies and SRA read data sets (genomic DNA and RNAseq transcriptome when available). Our analyses also involved coding of intron presence/absence and intron phase (codon phase) with an intron-coding scheme that was designed specifically for this study (detailed below). A robust scoring system for donor site motifs (5’-intron splice sites) was used based on the insect IUPAC motif “R|<u>GTRAGT…</u>” where “|” = the beginning of the intron (underlined). In this scoring system, each position that matches exact nucleotides in the motif “R|**<u>GT</u>**R**<u>AG</u>**T” contribute 1.0 points to the overall score (6.0 points total possible), while transition-type differences from these positions receive 0.5 points and transversion-type differences receive no points. Matches to degenerate positions in the motif receive 0.5 points at most due to degeneracy “**<u>`R`</u>**|GT**<u>R</u>**AGT”. The acceptor motifs (3’-intron splice sites) were also annotated to generally assess the quality of the splice sites, but these were not generally scored. The annotated acceptor motifs were annotated together with branch acceptor site motifs, which typically lie immediately upstream of acceptor sites. The combined branch site/acceptor motif used was “`TTYTNAY`…`Y`_9_`YAG|R`”.

### Intron-coding scheme

To facilitate data collection and curation, we implemented a novel intron-codification scheme that could be used in FASTA formatted protein sequences. Our intron codification encodes both the intron position and the intron phase with a modified or extended single-letter amino acid coding system. Each exon of a gene begins on a new line and starts either with the starting methionine (“M”) or the intron-encoding in lower case. Intron encodings are written in lower case and begin with an “`x`”, followed by a series of 3–5 “`w`” characters and ending in an “`x`”, followed by true amino acid sequence in uppercase letters. As intron codes only occur at the beginning of a line and are the only sequence characters using lower-case letters, they are easy to remove with a simple RegEx substitution command. Intron codes feature 3-5 “`w`” characters in between flanking “`x`” characters, which serve to highlight an intron code when a multiple sequence alignment (MSA) is visualized on the screen. The number of “`w`” characters in an intron code signify phase 0 (“`^xwwwx`”), phase 1 (“`^x`wwwwx”), and phase 2 (“`^x`wwwwwx”) introns, where “^” represents the beginning of a line (i.e., a RegEx positional anchor character). The choice of “`w`” was made because tryptophan residues (W) are the least common amino acids. Intron-coding sequences were left in place for sequence alignment but were removed prior to phylogenetic analysis of an MSA. The only exception was for early testing of the effects of our intron-coding scheme on alignment. This led to an expansion of the intron-coding scheme to allow the coding of intron absences, which is useful when the intron-coding scheme affected alignment. Intron absences were coded only when necessary in pre-trimmed FASTA files and never in our full-length FASTA files. Intron absences were encoded by three “`f`” characters (“^x<u>fff</u>x”, “^x<u>fff</u>xx”, and “^x<u>fff</u>xxx”) featuring 1–3 ending “`x`” characters to equal the width of the intron code at that position in other sequences. Last, “`y`” characters were occasionally used in place of “`w`” characters in the pre-trimmed FASTA files to indicate a derived phase change in an intron, an event that is quite rare in this data set. Overall, this new intron-coding scheme allowed us to identify errors in previous sequences subjected to automatic gene annotation. Fig. S7 demonstrates this intron coding for the fourth *Tmc123* intron, which changes phase in Pipunculidae and Schizophora.

### Sequence alignment

Sequence alignment was done using MUSCLE (54) on MEGA12 (55) with a relaxed gap insertion penalty of -2, which was possible because the intron-coding scheme constrained alignment of amino acid sequences encoded by homologous exons. Sequence alignment also involved pre-trimming. Pre-trimming of tandem repeats, novel exitrons, or lengthy loop insertions was done on “pre-trimmed” FASTA file versions of the “full-length” FASTA files. Pre-trimming was done by introducing a line-break in the FASTA file right before the desired trimming. For example, if an amino acid sequence featured 5 consecutive glutamine residues (“…`QQQQQ`…”) that we wished to trim down to three residues, we would introduce a line break right before the repeating Q’s, which would then be trimmed down to three Q’s at the beginning of the next line (“`^QQQ`…”). This allowed us to easily track and sometimes reverse pre-trimming choices. Thus, pre-trimmed FASTA files feature additional line breaks besides those used for intron-coding.

### Phylogenetic analyses

Final phylogenetic analyses were conducted using maximum likelihood (ML) estimation, and either a Jones-Taylor-Thornton (56) or a Le-Gascuel (57) amino acid substitution model with a gamma distribution with invariant sites (F+G+I) with 500 to 1000 bootstrap replicates on MEGA12. Both JTT and LG were indicated as the top two models under explicit model tests and performed similarly in different ML runs. A partial data cut-off from 15% to 30% was used for each MSA (explicitly stated in each tree figure along with the resulting number of alignment columns). For the CIB draft trees, more distantly related EF-hand containing proteins outside of the CIB family were used to verify that curated genes encoded one of the three main CIB genes and not a distantly related paralog (draft trees with distant paralogs not shown).

## Acknowledgements

This work was supported by an NSF-funded Evolutionary Sciences REU fellowship to C.B. (NSF DBI Award #2149361), and a 2026 summer support award from the University of Iowa’s Dept. of Biology to H.R. The authors would like to thank Andrew Forbes, Daniel Eberl, and Tina Tootle for comments on earlier versions of this manuscript.

## Contributions

A.E. conceived and led the study. H.R., C.B, and A.E. conducted all gene annotation and curation. A.E. and C.B. devised the intron-encoding scheme. A.E. conducted sequence alignments and phylogenetic analyses. A.E. and J.L. analyzed and discussed all phylogenetic results. A.E. and J.L. calculated the lower-bound estimates of gene losses. A.E. and H.R. constructed the figures and prepared the final submission. A.E. wrote the manuscript and all authors read and edited the final submission.

## Ethics declarations

Competing interests: The authors declare no competing interests.

---

**Fig. S1.**
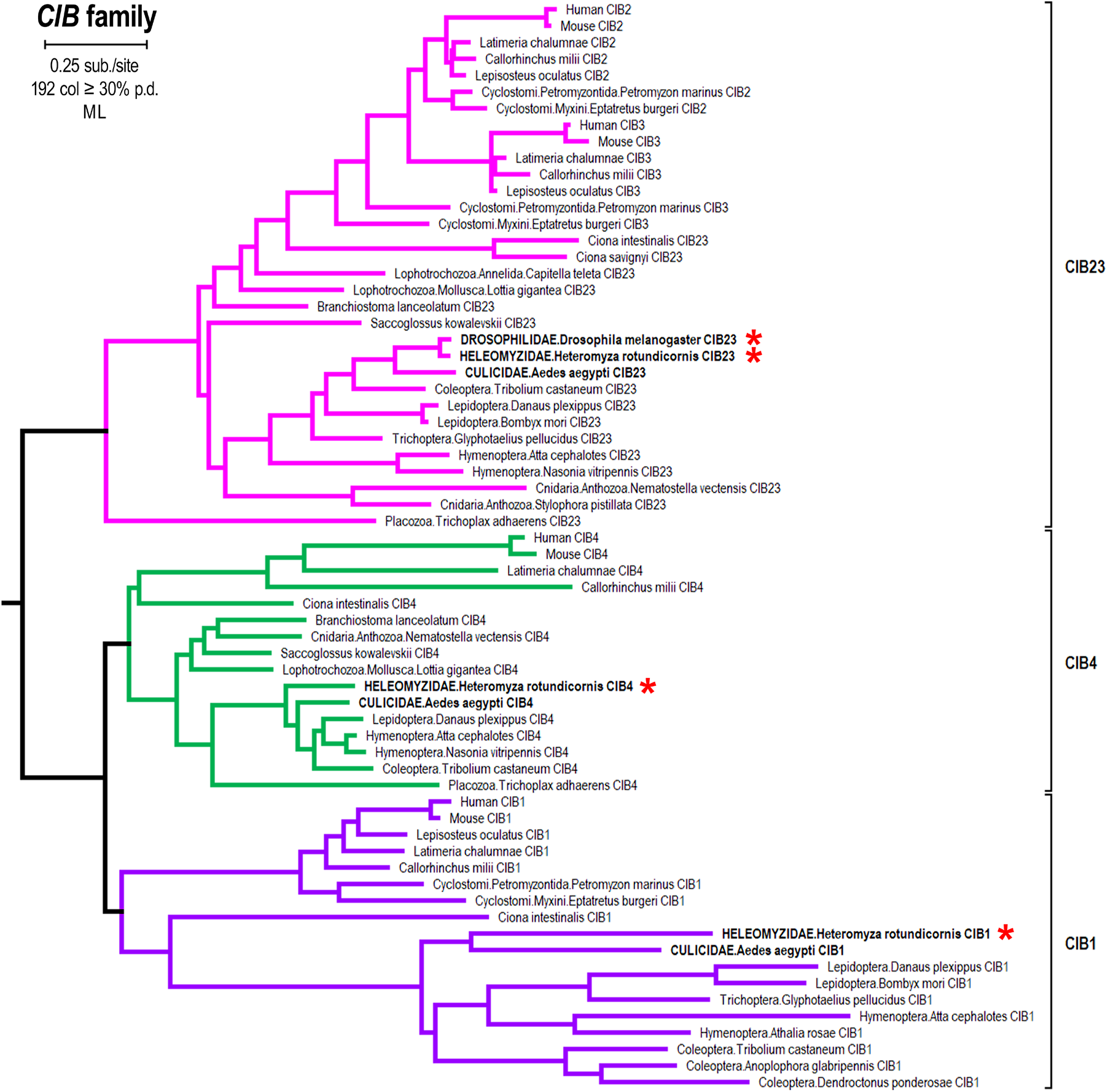
Phylogenetic tree of eumetazoan CIB genes. Like the eumetazoan TMC phylogeny (see Fig. 1), eumetazoans have three CIB genes: *CIB23* (magenta clade), *CIB4* (green clade), and *CIB1* (purple clade). Also, like the eumetazoan TMC phylogeny, *Drosophila melanogaster* (but not *Heteromyza rotundicornis*, a basal schizophoran fly) is missing *CIB4* and *CIB1* (compare red asterisks). Unlike the TMC phylogeny, eumetazoan CIB genes are not as duplicated in jawed vertebrates; only *CIB23* duplicated into *CIB2* and *CIB3* in a stem-vertebrate ancestor. Because the CIB genes are short protein-coding sequences (∼600 bp) and because *CIB4* and *CIB23* have many invariant sites, this phylogeny does not ideally recapitulate relationships within each paralogy clade. Nonetheless, there is sufficient phylogenetic signal in this data set, including paralog-specific peptide motifs and introns, to confidently assign membership to one of the three paralogy clades.

**Fig. S2.**
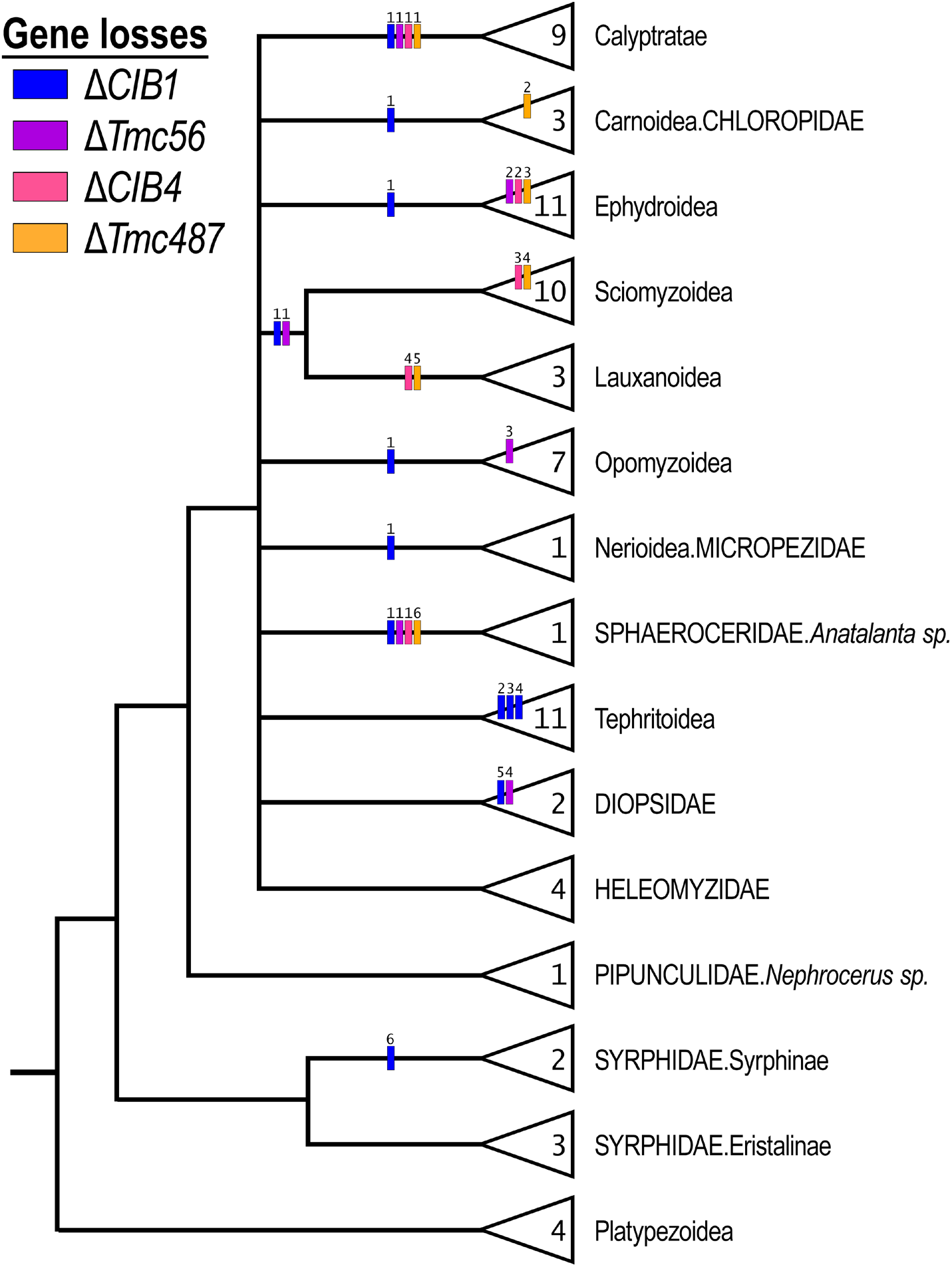
Lower bound estimates of TMC and CIB gene losses in Cyclorrhapha. Shown is a cladogram of Cyclorrhapha with Schizophora conservatively collapsed into a 10-way polytomy to estimate the minimum number of gene losses. Gene loss characters are color-coded according to the key in the upper left-hand corner and indexed within the tree. Loss characters sharing the same index number are counted as a single character. Loss characters within the labeled clades (triangles) occur within a specific lineage of that clade. The number of taxa analyzed in each clade is listed in each triangle. Altogether, the TMC and CIB data set for Cyclorrhapha requires a minimum of 4–6 separate gene losses for each of gene for a lower bound estimate of 19 separate gene losses. This lower bound does not include pseudogenes as losses. See Table 1 for a lower bound in which pseudogenes are counted as losses.

**Fig. S3.**
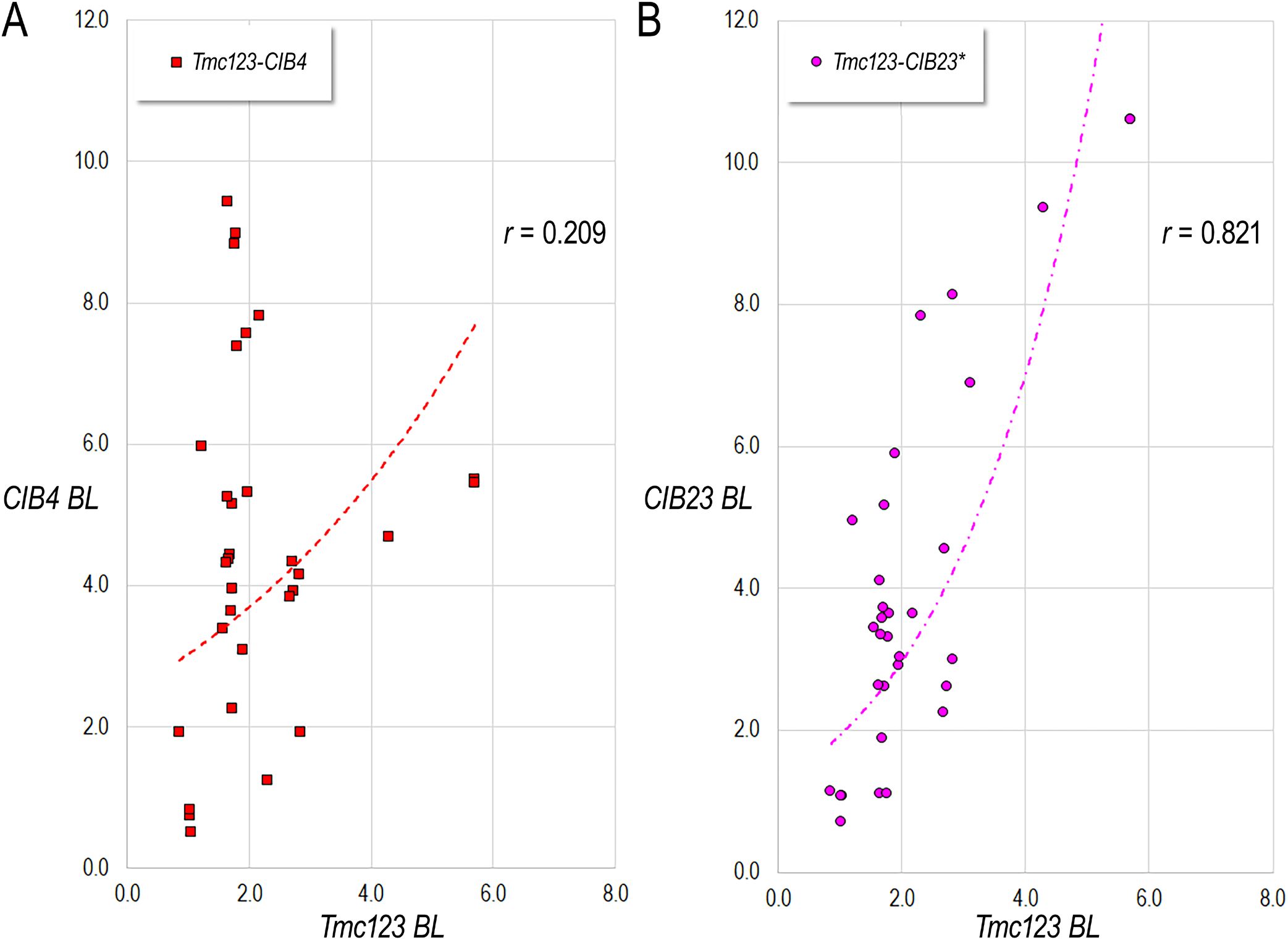
Mismatched *TMC-CIB* branch lengths are uncorrelated. **(A)** Shown is a control scatterplot between *Tmc123* and *CIB4* BLs for lineages that have *CIB4* genes. Compared to the *Tmc123-CIB23*, *Tmc487-CIB4*, and *Tmc56-CIB1* pairings shown in the Figure 3B scatterplot, this *Tmc123-CIB4* comparison is not tightly correlated. Trendlines here are exponential fits. Pearson’s correlation coefficient (*r*) is given in the upper right-hand corner. **(B)** For comparison, shown is a scatterplot between *Tmc123* and *CIB23* BLs for lineages that have *CIB4* genes (*Tmc123-CIB23*\*) shows that these are highly correlated (*r* = 0.821) unlike the low correlation of *Tmc123-CIB4* (*r* = 0.209).

**Fig. S4.**
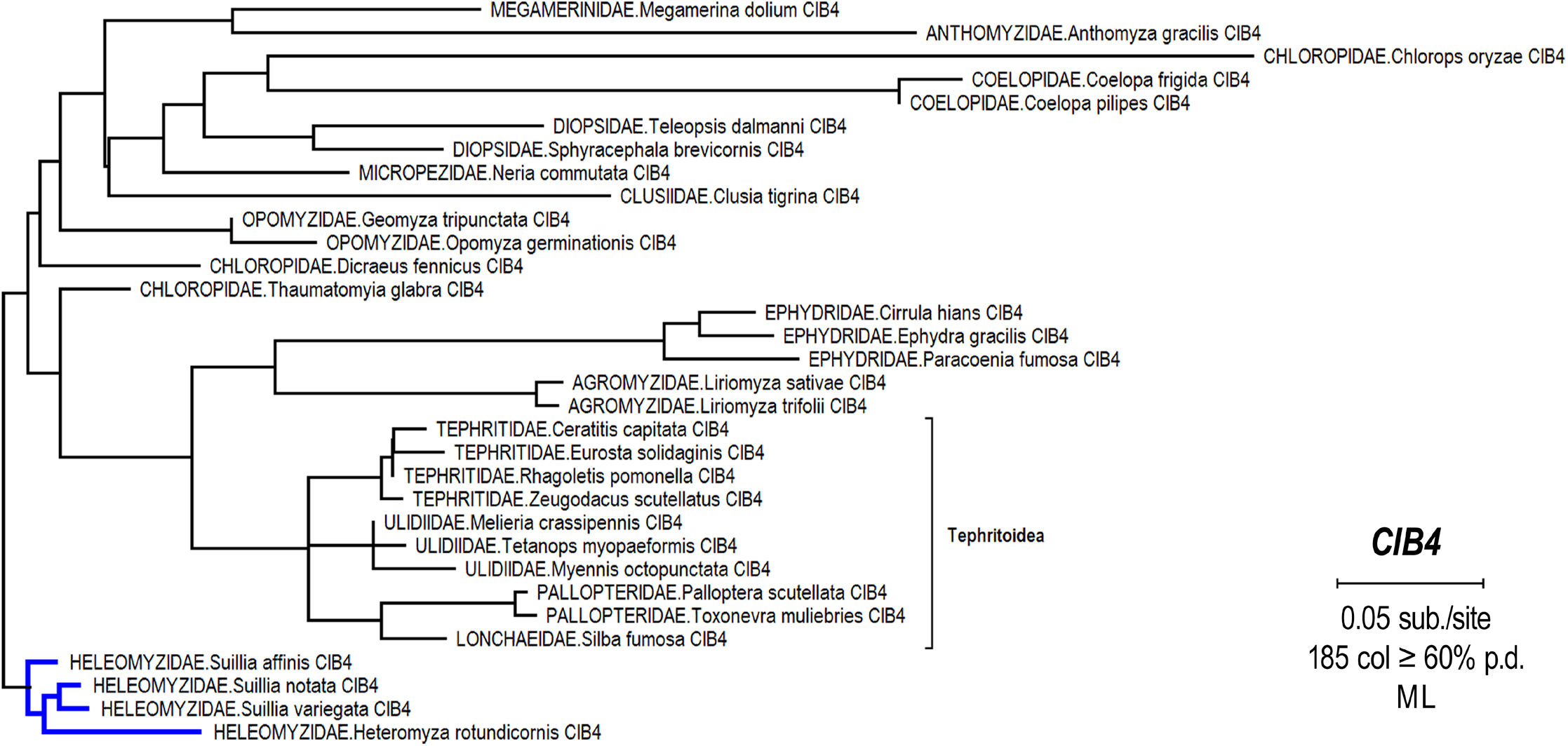
Phylogenetic tree of schizophoran *CIB4*. Maximum likelihood estimation was conducted on all available schizophoran *CIB4* sequences to date and rooted with Heleomyzidae as outgroup (blue clade). Note that this tree is somewhat distorted due to inclusion of fast-evolving (long-branched) sequences and because this is a slow-evolving gene (compare scale bar to other trees in this study). This tree was used to compute heleomyzid-normalized branch lengths.

**Fig. S5.**
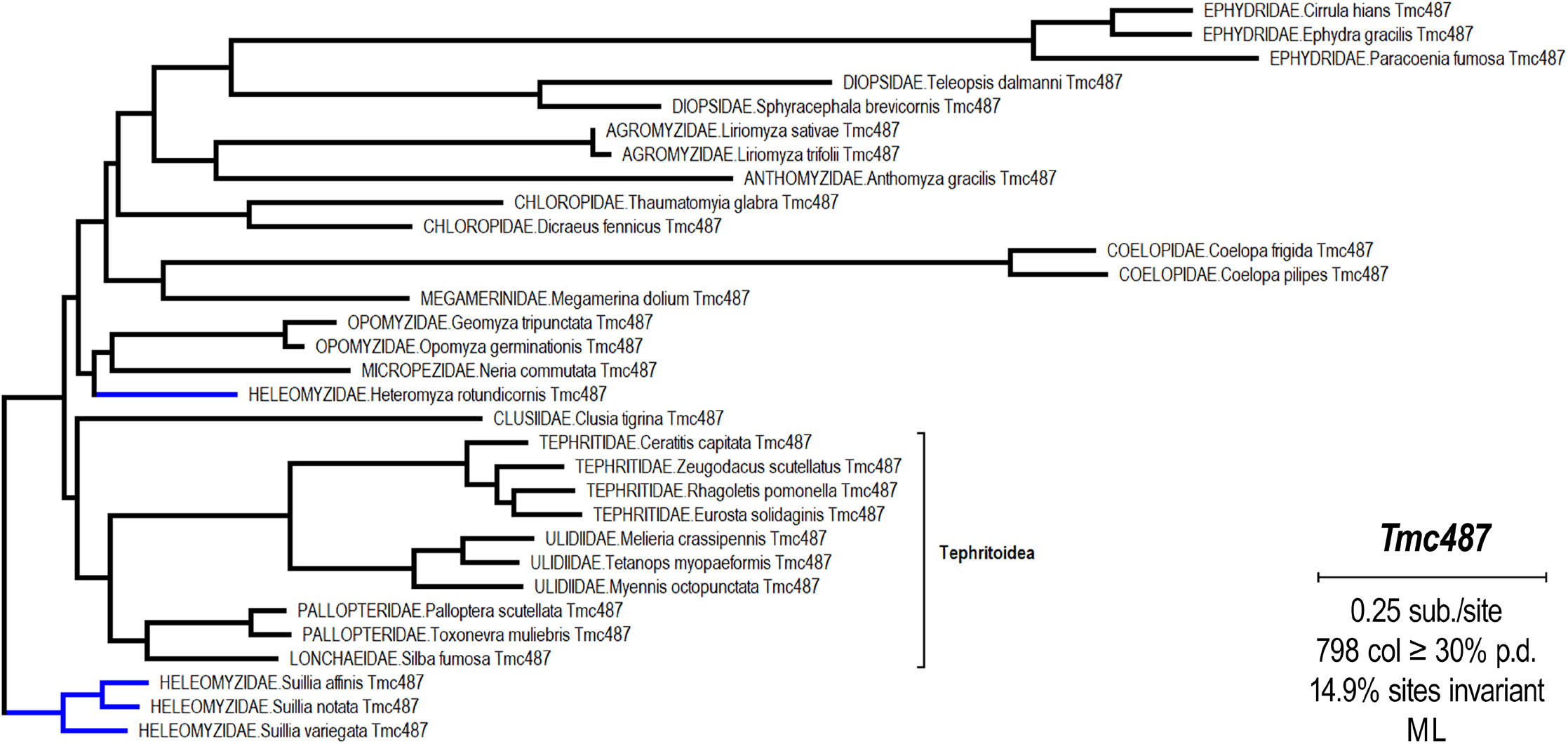
Phylogenetic tree of schizophoran *Tmc487*. Maximum likelihood estimation was conducted on all available schizophoran *Tmc487* sequences to date and rooted with the bulk of Heleomyzidae (*Suillia* genus) as outgroup (blue lineages = heleomyzids). Note that this tree is somewhat distorted due to inclusion of fast-evolving (long-branched) sequences. This tree was used to compute heleomyzid-normalized branch lengths.

**Fig. S6.**
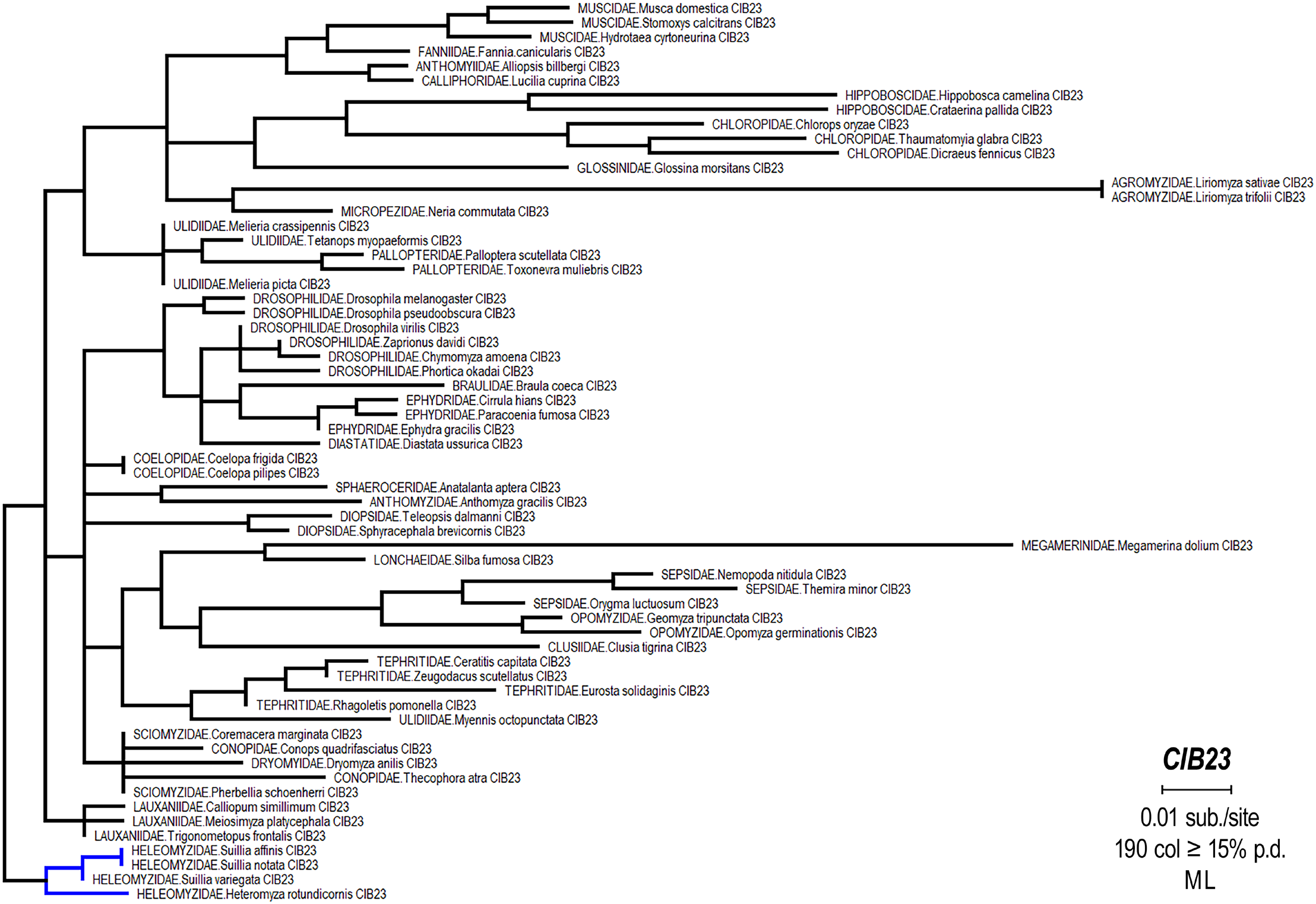
Phylogenetic tree of schizophoran *CIB23*. Maximum likelihood estimation was conducted on schizophoran *CIB23* sequences and rooted with Heleomyzidae as outgroup (blue clade). Note that this tree is somewhat distorted within the in- group because this is a slow-evolving gene and a few independent changes result in long-branch attraction (compare this tree’s scale bar to the scale of other trees in this study). This tree was used to compute heleomyzid-normalized branch lengths.

**Fig. S7.**
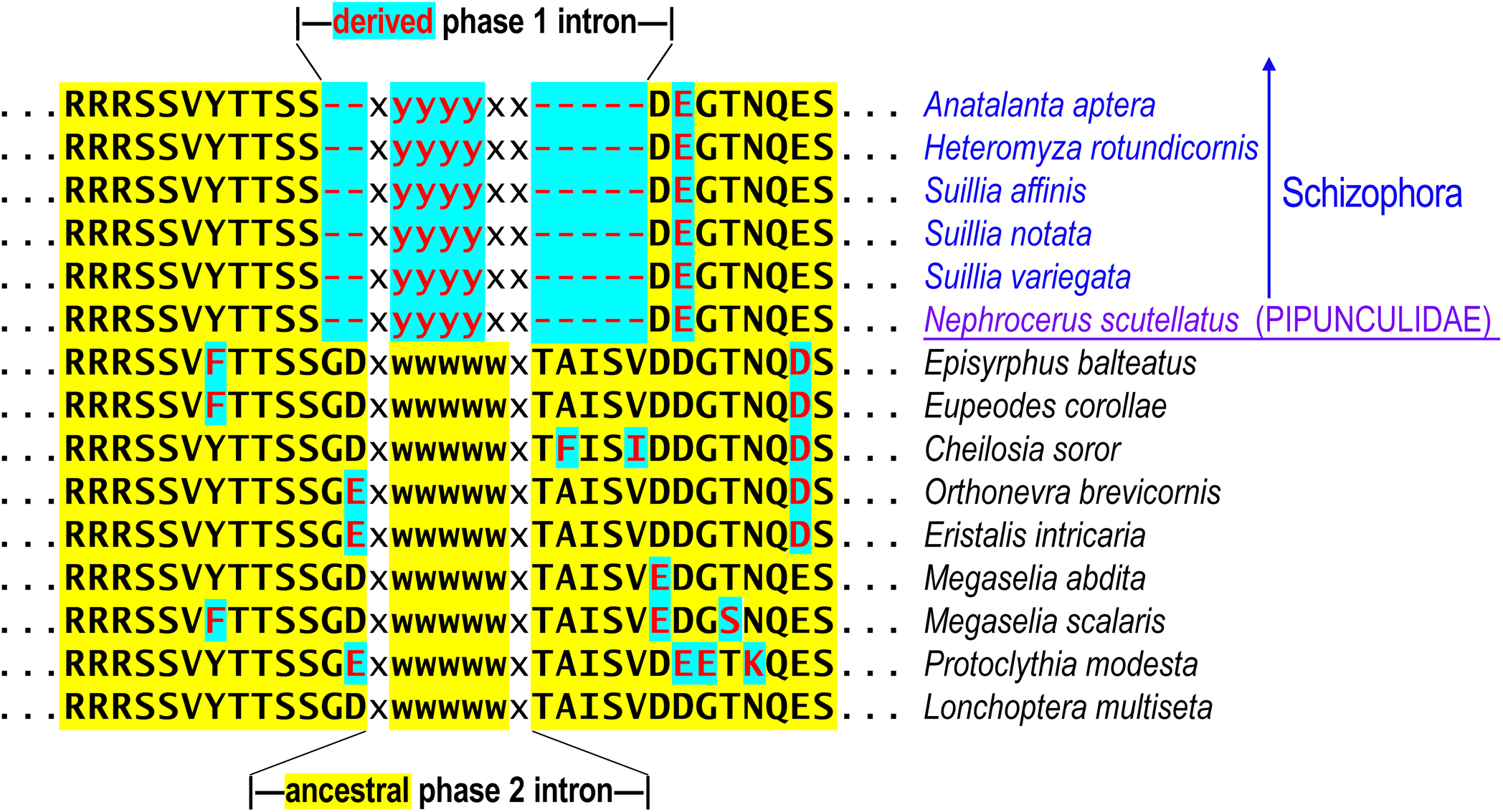
A molecular synapomorphy uniting Schizophora and Pipunculidae. Shown is the amino acid sequence of the *Tmc123* flanking intron 4 encoding (phase two “xwwwwwx” and derived phase one “xyyyyxx”). Intron 4 of *Tmc123* was ancestrally a phase two intron (basal cyclorrhaphans listed below *Nephrocerus scutellatus*), which changed to a phase one intron in the lineage leading to both Pipunculidae (big-headed flies such as *N. scutellatus* underlined below) and Schizophora (representative sequences from Sphaeroceroidea above *N. scutellatus*). This change is associated with a deletion of 7 amino acids flanking both sides of the intron. This is an example of a molecular synapomorphy (a shared derived trait). Intron phase changes are quite rare in the TMC and CIB data sets relative to intron gains and losses. Yellow-highlighted text corresponds to conserved ancestral characters, while cyan-highlighted red text signifies derived changes. See Methods for a more detailed explanation of the intron-encoding scheme.

